# Single-cell phenotypic characteristics of phenotypic resistance under recurring antibiotic exposure in *Escherichia coli*

**DOI:** 10.1101/2021.05.26.445729

**Authors:** Silvia Kollerová, Lionel Jouvet, Julia Smelková, Sara Zunk-Parras, Alexandro Rodríguez-Rojas, Ulrich K. Steiner

**Affiliations:** Department of Biology, University of Southern Denmark, Odense, Denmark; Biological Institute, Freie Universität Berlin, Berlin, Germany

**Keywords:** single-cell, microfluidics, antibiotic resistance, β-lactam antibiotics, growth, survival, persistence, tolerance

## Abstract

Despite increasing interest, non-heritable, phenotypic drug resistance, such as tolerance and persistence towards antibiotics, remains less characterized compared to genetic resistance. Non-heritable drug resistance challenges antibiotic treatment and have implications towards heritable resistance evolution. Phenotypically resistant cells have commonly been characterized as growth arrested cells prior and during antibiotic application that quickly resume growth post-application. Here, we add novel combinations of characteristics of phenotypic resistant *E. coli* single cells—that are of particular interest towards genetically fixed resistance—, and contrast those to susceptible cells of the isoclonal initial population by exposure to different levels of recurrent antibiotic. We find that phenotypic resistant cells reduced their growth rate by about 50% compared to growth rates prior to antibiotic exposure, but cells do not go into near growth arrest. The growth reduction is induced by antibiotic exposure and not caused by a stochastic switch or predetermined state as frequently described. Cells exhibiting constant intermediate growth survived best under antibiotic exposure and, contrary to expectations, selection did not primarily act on fast growing cells. Our findings support diverse modes of phenotypic resistance, and we revealed resistant cell characteristics that supports acclaims of an underappreciated role of phenotypic resistant cells towards resistance evolution.

## Introduction

Resistance to antibiotics, in combination with slow progress in discovering novel antibiotics, poses a major challenge to modern medicine (1). Antibiotic resistant cells sustain bactericidal concentrations of antibiotics through heritable, often well-known and explored, genetically fixed mutations. In contrast, non-heritable phenotypic resistant cells, also often referred to as antibiotic persistent or tolerant cells are in a non-heritable drug resistant phenotypic state. This state allows them to survive periods of bactericidal concentration of antibiotics and is classically characterized by non- or very slow growth prior and during exposure to the drug (2–5). More recently ‘specialized’ persister cells have been described that show less growth dependent persistence (6–12). Identifying persister cells is usually done through a characteristic biphasic killing curve that is observed at the population level and caused by differential selection of persister cells of a susceptible population during antibiotic exposure (2,13), in contrast tolerant cells do not show a biphasic killing curve as they are assumed to be a population wide response and not a subpopulation one (2–5,9). However, definitions on persistence and tolerance are not consistent as persistence is also referred to as a special case of tolerance, or is defined as hetero-tolerance (4). The persister cell stage can be environmentally induced or stochastically switched to, hence, cells can dynamically alter states, and progeny of persiter cells are as susceptible as the parental population from which they derived (4,14). Quantitatively, below 0.1% of cells in wild-type susceptible populations are persister cells, though variation in persister fractions exists among experiments, strains, growth media, and environmental conditions (15). Uncertainty in accurately quantifying persister cell fractions based on killing curves comes from highly sensitive determination of slopes and intercepts of killing curves (10).

Tolerance is a population level response and leads to a slowed killing curve, tolerant cells are therefore much more common than persister cells (2,9,16). Tolerant cells tolerate antibiotic exposures by mainly non- or very slow growing under antibiotic exposure, characteristics they share with persister cells (2). Though compared to persister cells, tolerant cells can only sustain shorter durations of antibiotic exposure (4). Under antibiotic application, tolerant cell populations show prolonged minimum duration of killing (MDK) without a change in the minimum inhibitory concentration (MIC), that is, they die slowly under antibiotic application even when exposed to very high concentrations.

A third class of cells that can sustain bactericidal concentration of antibiotics by phenotypic resistance are viable but non-culturable cells (VBNC/ deep dormant cells / sleeper cells) (Dong et al 2020). In contrast to persister and tolerant cells, VBNC cells do not regain growth 3-4 hours post-antibiotic exposure, but instead remain at low metabolic activity in a deeper dormancy state and do not easily resuscitate, potentially due to a lack of intracellular ATP (17). Persister and tolerant cells are therefore more susceptible to recurring exposure to antibiotics compared to VBNC cells (17). VBNC cells are further phenotypically characterized by being small (2.3 ± 0.2μm for *E. coli*) and less elongated compared to persister cells (18).

Differentiating among these different phenotypic resistant cell types can be difficult, in particular when single-cell data is not available (3,4,7,19). A growing number of studies on persistence work at the single-cell level and primarily aim at a mechanistic understanding of phenotypic resistance (8,18,20–24), yet in depth analyses of single-cell phenotypic characteristics of phenotypic resistant cells are still rare (8,10,25). Many of these single-cell studies use persister mutants, and start with growth-arrested, stationary phase cells, as these conditions increase the fraction of persister cells. Some recent studies suggest that persister cells arising of exponential growing populations are predominantly not growth arrested prior to antibiotic exposure (10). Despite evidence for diverse modes of phenotypic resistance, overall, these single-cell level studies highlight that that persister cells show limited growth when exposed to β-lactam or other antibiotics that inhibit cell wall formation or cell division (6,7,26), and that switching to persister stages can be irreversible or a reversible dynamic stochastic process (6,10). Earlier studies reported that the stochastic switch to the rare persister state occurred for growth-limited cells within exponentially growing *E. coli* populations, with ~80% of persisters arising from the non-growing sub-population prior to treatment, while the remaining persister cells originated from cells that were growing prior to treatment (11,27). Other high throughput experiments, not working on persister mutant strains, suggest that large fractions of persister cells arise from non-dormant cells when selected of an exponentially growing population (10). Quantitative high-quality single-cell data on phenotypic characteristics of phenotypic resistant cells are limited, as persister cells are so rare and selecting for them remains challenging (10). However, characterizing single-cell growth, division and mortality of phenotypically resistant cells would provide deeper understanding about the evolutionary importance of non-heritable resistant cells for the evolution of heritable (genetic) resistance and efficacy of antibiotic treatment, as high frequencies of non-heritable tolerant cells provide evolutionary reservoirs and increase the probability of genetically fixed resistance mutations to occur (28–37).

Here, we use a demographic and phenomenological approach to quantify the fraction of cells in a phenotypic resistant stage among susceptible stage cells, we investigate how phenotypic resistant cells differ in their single-cell characteristics compared to susceptible cells, and we study what triggers changes in their phenotypes. To increase the number of cells that sustained antibiotic exposure, we reduce exposure periods to recurring 1.5 h periods compared to a typical single 3-5h exposure period. Whether the resulting selected cells should be referred to as hetero-tolerant cells, or persiter cells depends on definitions, we follow with the term persister. We started with a susceptible clonal exponentially growing population of cells that we expose to recurrent antibiotic treatment at sub- to supra level antibiotic concentrations, we track each cell for their growth, division, and survival in a microfluidic device called a mother machine (38–40). By exposing cells to a β-lactam, ampicillin, that corrupts cell wall synthesis, we expected quickly-growing cells to be heavily selected against, leaving primarily slow-growing phenotypic resistant cells that also arose from the slowest-growing population of cells prior to antibiotic exposure (3). Phenotypic resistant cells, compared to genetically resistant cells, should not be sensitive to antibiotic concentrations beyond the MIC of the susceptible cell population (3), and are expected to regain normal growth rates within 3-4 hours post-antibiotic treatment, as long as they are not VBNC cells. We expected few cells to stochastically switch to tolerant resistance stages or switch by environmental induction, as we expected phenotypic resistant stages to be mainly predetermined prior to antibiotic exposure. To evaluate how our findings at the single-cell level might scale to cell culture dynamics, we exposed cell cultures to similar recurring exposures of antibiotics and counted colony forming units (CFU) before and after each exposure to antibiotics. In addition, we performed an experiment by measuring the optical density dynamics of constantly exposed cell cultures to the same levels of antibiotics. We discuss why scaling and quantitative comparison between single-cell characteristics and experiments, and cell culture level exploration can be challenging.

## MATERIAL METHODS

We conducted four set of experiments, three of them with three recurring antibiotic (ampicillin) exposure periods of 90 minutes; exposure periods started 60 min, 330 min and 1080 min past onset of the experiments. Two of these sets collected single-cell data using a mother machine microfluidic device and time-lapse microscopy (Movie SI) (38,39,41) and one set collected cell culture level data, i.e., CFU counts (details SI). Cells were derived of an exponentially growing susceptible isoclonal *E. coli* strain K12-MG1655, ATCC® 700926TM, population. Exposure was at one of eight different concentrations ranging from sub-MIC to supra-MIC levels (0-128μg/ml; details SI). Cell death in the microfluidic device was determined using propidium iodide. The fourth experiment, which was at the cell culture level, continuously exposed cells to antibiotics and served as a control experiment. All experiments, single-celled and cell culture experiments, were done in supplemented minimal media (M9), and all cultures were derived from the same *E. coli* strain, that have a replication time of ~25-30 minutes in M9 medium at 37°C (details SI). We chose this strain as it is well adapted and explored and has been used in previous work on persister cells (11,18,26,42). More details on bacterial cell culturing, microchip casting and mounting, microchip loading, life cell imaging, image analysis, experimental treatment are provided in the SI. We remain brief in our description here as most methods follow previously published protocols (39,43).

The resulting data was subsequently analysed in program R (44) using general linear, generalized linear, and nonlinear (Generalized Additive Models, GAM) models. Models were compared based on information criteria (AIC)(45). We considered an ΔAIC >2 as better support between competing models. Survival analyses (Kaplan-Meier Fig. 1a) are computed with R package survival (survfit function) and compared amongst them with a Cox proportional hazard model. Probability of death curves (Fig. 1b, Fig. 3b), division rate curves (Fig. 2a), and size curves (Fig. 2b) were plotted with ggplot2 package and a loess smoothing (geometric_smooth function; package ggplot2), growth curves were fitted with loess smoothed GAM models (Fig. 2c), competing GAM models (Table 1) are fitted with the restricted maximum likelihood method (REML). Details on the statistical testing, exclusion of extreme values, and model specifics are given in the SI, which also include details on comparisons of growth differences before death (GLM’s; post-hoc Tukey test using glht function of mutlcomp package), and testing for the robustness of growth rates (Fig. 4; GAM).

**Fig. 1:**
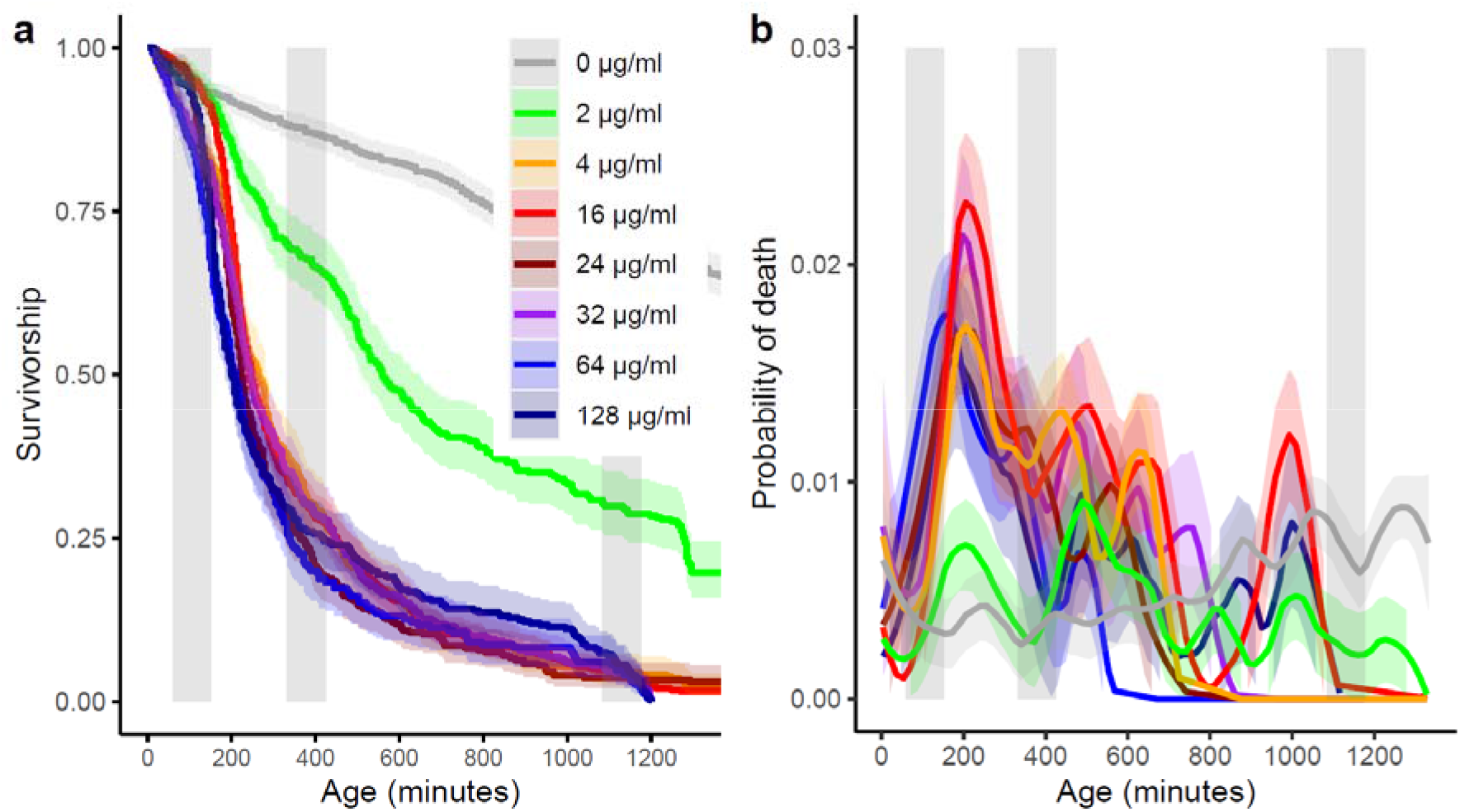
Survivorship curves (a) and probability of death, i.e., force of mortality (b) of cells exposed to different levels of antibiotics across the duration of the experiment. The grey curves are control group cells that were not exposed to any antibiotics. The grey vertical bars mark the exposure period to the antibiotic. Kaplan-Meier survivorship curves (a) are shown with 95% CI. Probability of death curves (b) are loess smoothed with 95% CI. Note, the two panels (a & b) are built from the same single-cell mortality data.

**Fig. 2:**
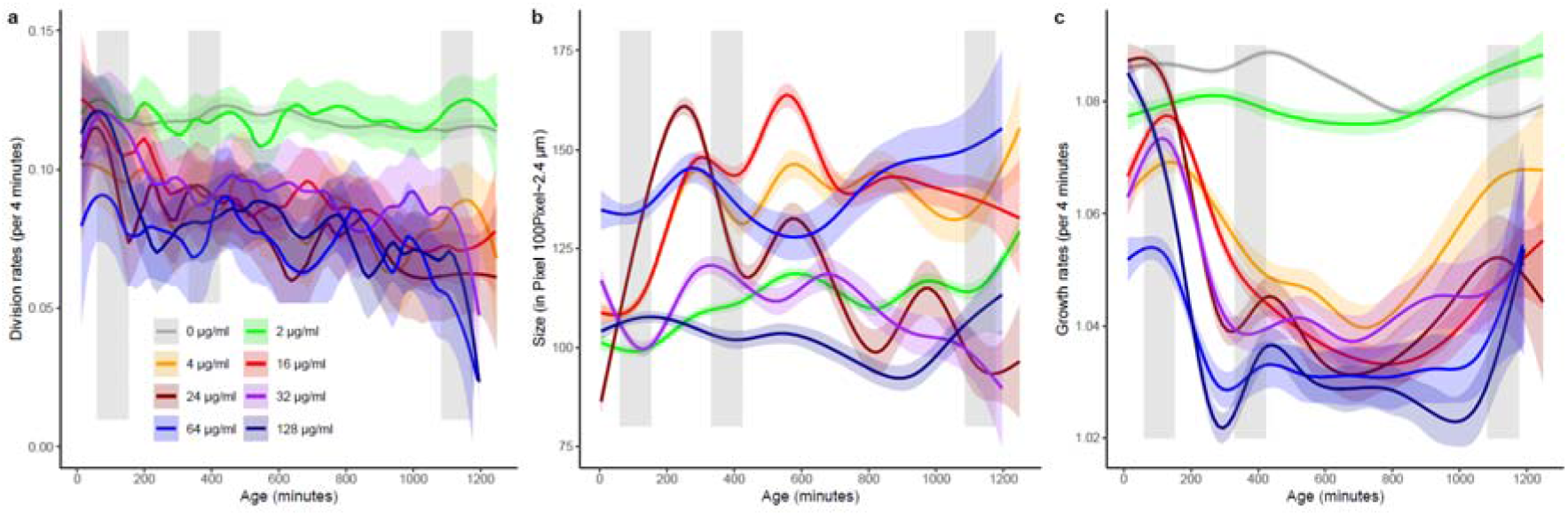
Cell division rates (a), cell size (in pixels with 100 pixels~2.4μm) (b), and growth rates (c) of cells exposed to different levels of antibiotics across the duration of the experiment. The grey vertical bars mark the exposure period to the antibiotics. Division rate (a) size (b) and growth rate (c) curves are with 95% CI. Note, size data (b) for the control group (0μg/ml) was not available.

**Fig. 3:**
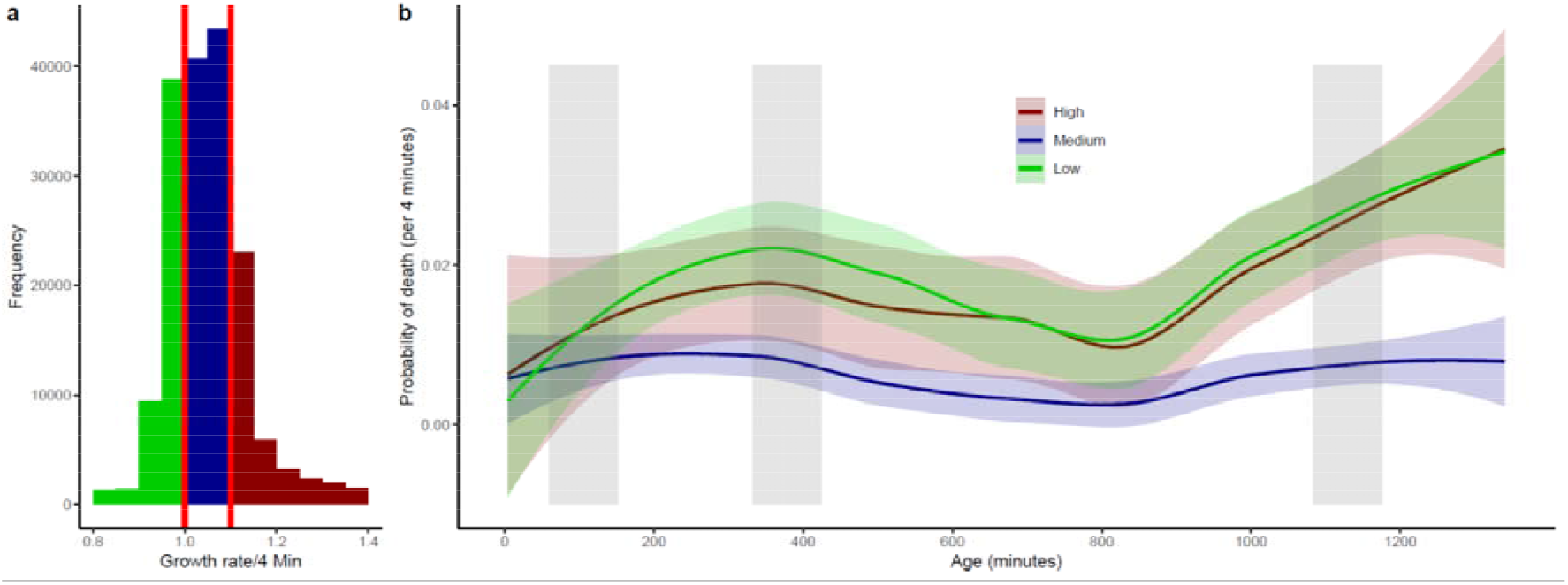
Cell growth rate distribution divided into three categories: growth arrested (shrinking or non-growing; green), intermediate growth (blue), and fast growth (red) (a), and probability of death for these three groups of growth arrested, intermediate, and quickly-growing cells across the duration of the experiment (b). Confidence intervals (95% CI) are shown in shading in panel (b). The grey vertical bars in panel (b) mark the exposure period to the antibiotics. Note that cells can switch among growth categories throughout the experiment, that is cells are categorized according to their current cell growth rate. For this analysis, we have combined cells across levels of antibiotic exposure ≥4 μg/ml and excluded all cells exposed to 2 μg/ml. A statistical model comparison showed that mortality differed substantially among the three growth category groups: ΔAIC 25.8 comparing a growth category model and a null model, see also Supplemental Information. Classical p-value post-hoc analyses showed that quickly growing and growth arrested groups did not differ from each other in age at death, but the intermediate group did.

**Fig. 4:**
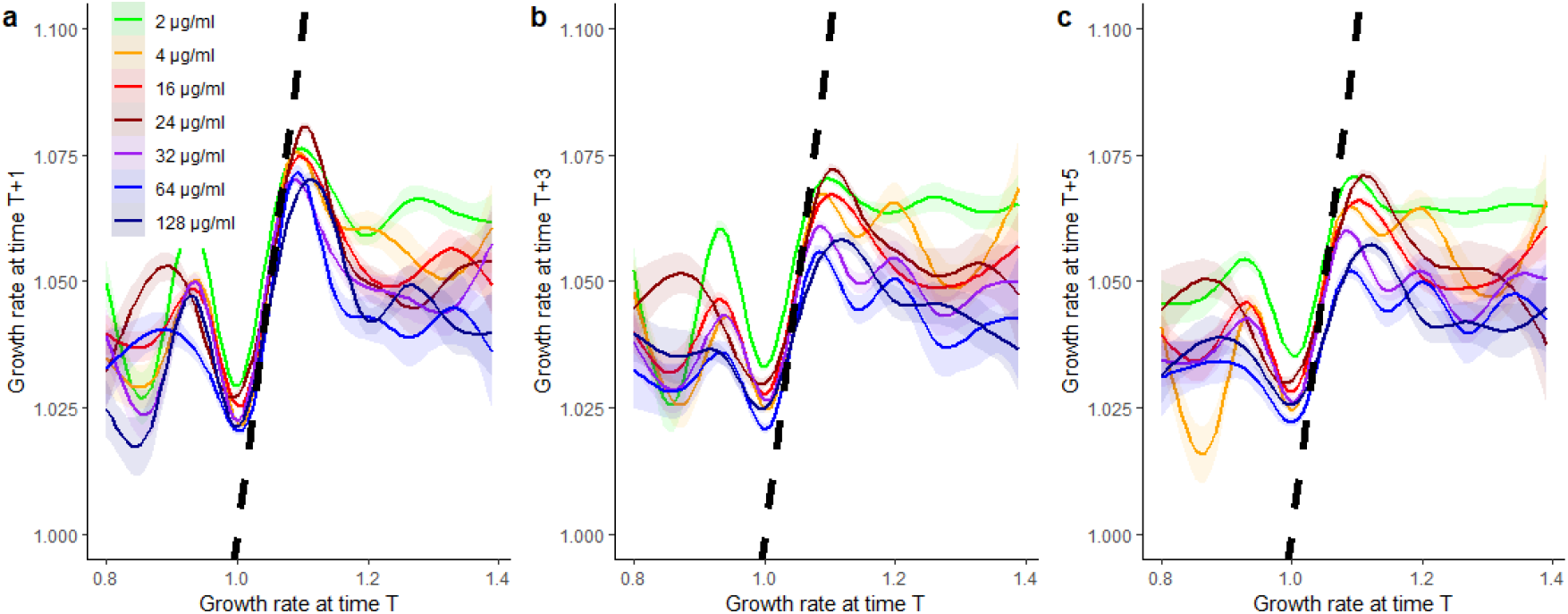
Correlation in cell growth rates at a given time t, compared to growth rates at time t+1 (4 minutes later) (a), t+3 (12 minutes later) (b), and t+5 (20 minutes later) separated out into populations recurrently exposed to different levels of antibiotics (as described in the main experiment). All growth rate correlations are fitted with a GAM with 95% CI. The dashed black line illustrates the direct correlation among time points.

**Table 1:**
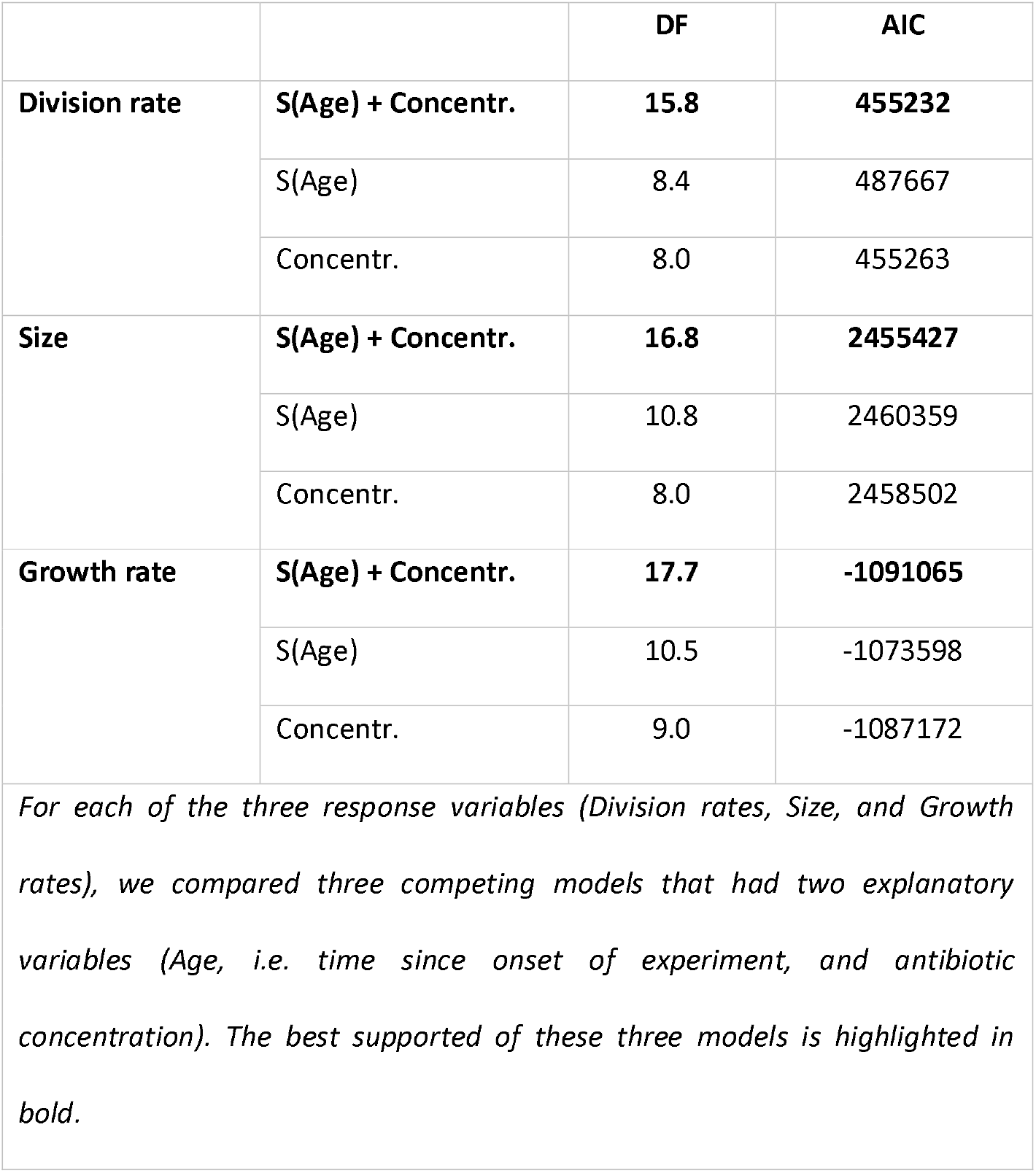
Model comparison among competing GAM models.

|  |  | DF | AIC |
| --- | --- | --- | --- |
| Division rate | **S(Age) + Concentr.** | **15.8** | **455232** |
|  | S(Age) | 8.4 | 487667 |
|  | Concentr. | 8.0 | 455263 |
| Size | **S(Age) + Concentr.** | **16.8** | **2455427** |
|  | S(Age) | 10.8 | 2460359 |
|  | Concentr. | 8.0 | 2458502 |
| Growth rate | **S(Age) + Concentr.** | **17.7** | **-1091065** |
|  | S(Age) | 10.5 | -1073598 |
|  | Concentr. | 9.0 | -1087172 |
| *For each of the three response variables (Division rates, Size, and Growth rates), we compared three competing models that had two explanatory variables (Age, i.e. time since onset of experiment, and antibiotic concentration). The best supported of these three models is highlighted in bold.* |  |  |  |

For the cell culture experiments, to collect the number of colony forming unit (CFU) data and the cell culture density data (OD_600_), we initially seeded these experiments using cells from an exponential growing culture. Each experiment had 8 replicates, the experiments were done in 96 well-plates, and cells grew at 37°C and vigorous shaking in their respective well (Supplementary File 2). Recurrent exposure to different antibiotic concentrations was achieved through serial dilution, centrifuging, removing super natant, resuspending and culturing cells, followed by serial dilution and plating on agar plates and CFU were counted after 16h of incubation, more details are provided in the SI.

## RESULTS

### Mortality and selection for tolerance

As expected, the probability of cell death, i.e., the force of mortality, increased after the first exposure of the susceptible parental population to the antibiotic, when exposed to MIC or higher concentrations (Fig. 1b, Movie SI). Selection for phenotypic resistant cells was strong with about two thirds to three quarter of cells dying before the second exposure period, when exposed to MIC or higher concentrations (Fig. 1a, Movie SI). The survivorship patterns (Fig. 1a) somewhat resemble a biphasic killing curve as frequently observed at the cell culture level of persister cells, though we highlight that direct comparison among cell culture level killing curves and single-cell survival can only be made in a crude way. Mortality peaked 30 to 60 minutes after the first 90-minute antibiotic exposure period, which suggests that the force of selection peaked with a lag time of 2 to 2.5 hours after the start of the first exposure. Survival patterns were similar at, or above, the MIC of 4 μg/ml (Fig. 1a, Cox proportional hazard model Likelihood ratio test 477.1, p<0.0001, n= 3496; for post hoc testing see SI Table S1). The subsequent 90-minute exposure period triggered a less pronounced mortality force compared to the first exposure period with a similar lag time of about 30 to 90 minutes after the second antibiotic exposure. Note, cell numbers became small (~20-30 cells per concentration) and estimates of mortality rates less reliable towards the third exposure period. For the control, 0 μg/ml, cells died, as expected, at a low rate and in agreement with experiments aiming at senescence patterns of aging single cells (40).

### Division rate, size, and growth responses

In contrast to the control group of no antibiotic exposure, cells that had been exposed to MIC or higher antibiotic concentrations reduced their division rate after the first exposure period, changing from ~2 to ~1 division per hour (Fig. 2a, Table 1). Most of this reduction in reproductive rate was reached ~150-200 minutes after the onset of the first exposure (300-360 minutes after onset of experiment). The division rates tended to only slowly decline after the second exposure, patterns that might also arise due to senescence and not only antibiotic exposure (41).

Average cell size remained approximately constant within a given antibiotic concentration. Although we found a surprising amount of variation in size among cells exposed to different levels of antibiotics (Fig. 2b, Table 1), there was no relationship between size and the levels of antibiotics. Cell growth rates responded as predicted by declining during and after the first antibiotic exposure for cells exposed to the MIC or above. We highlight here that growth rates only approximately halved (Fig. 2c, Table 1), which contrasts our predictions based on previous findings where persister cells were growth arrested or near growth arrested (4). The induced growth reduction started about 30 minutes after the onset of antibiotic exposure, which is a lag time that corresponds to approximately the doubling rate (division time) of the parental population. The lowered growth rates stabilized about 4-5 hours after onset of the initial exposure period, suggesting substantial variance in lag time among individual cells’ growth response. Induced growth reductions depended partly on antibiotic concentrations with higher levels of antibiotics leading to lower growth rates. However, even with the highest antibiotic concentration, the growth rates did not drop below 1.02-1.03/4 minutes, i.e., 2-3% cell elongation within 4 minutes. Further, growth did not resume to prior exposure levels rates within 3-4 hours after release of exposure.

### Growth specific selection

The growth rates as discussed above (Fig. 2c) are average growth rates at a given point in time, and even though variance in growth rates (see CI Fig. 2c) does not increase substantially during the experiment (beyond the expected increase due to decreasing number of cells towards the end of the experiment), selection might still act preferentially against cells with high growth as expected from a β-lactam antibiotic and as previously described for phenotypic resistant cells (14). To evaluate the possibility of such differential selection based on cell growth we analysed together all cells exposed to MIC or above, and then separated them out into three growth categories: cells that did not grow or shrank (growth arrested cells with growth rates <1.0), cells that grew at low to intermediate rates (growth rates 1.0-1.1), and cells that grew quickly (growth rates >1.1) (Fig. 3a). Note that cells can switch dynamically among these categories throughout the experiment and that mortality and survival is associated only with the current growth rate of a cell and not a cell’s growth rate averaged across time. As expected, quickly growing cells showed increased probabilities of death upon exposure to antibiotics, but contrasting to our expectations, the probability of death of growth arrested cells increased substantially upon exposure to antibiotics, whereas the probability of death of slow-intermediate growing cells did not change (Fig. 3b, see also SI Table S2 and Supplementary file 1C).

To verify the robustness of our finding that selective mortality is dependent on growth rates, and to broaden our understanding of how growth influences mortality beyond the current growth rate (Fig. 3b), we did three comparisons between average growth rates of cells dying during specific periods in time. First, we compared the average growth rate (averaged across time) of cells that survived the first exposure period to those that died before the first exposure period. Second, we compared the average growth rate of cells dying during or shortly after the first exposure period (cut off at peak of first mortality) to those surviving past the first peak of mortality, and third we compared similarly for cells that died between the first peak of mortality and the second peak of mortality to those that survived past the second peak of mortality (Table 2). Before the first exposure, cells that died had lower average growth rates compared to those that survived past the first exposure, showing that without selection of antibiotics slower growing cells are preferentially selected against. Cells that died during the first exposure or shortly after, i.e., during times of strong selection, had slightly higher growth rates than those that survived past this first increased selection period. This pattern of growth dependent selection remained for the next period of increased selection after the second antibiotic exposure. These findings support our initial prediction that faster growing cells would be preferentially selected against under antibiotic exposure. However, note that the effect sizes (growth differences) among surviving and dying cells are small, even though they are significantly different. Combined with the findings shown in Fig. 3, we interpret our findings on growth specific selection under antibiotic exposure as not primarily acting on fast-growing cells.

**Table 2:**
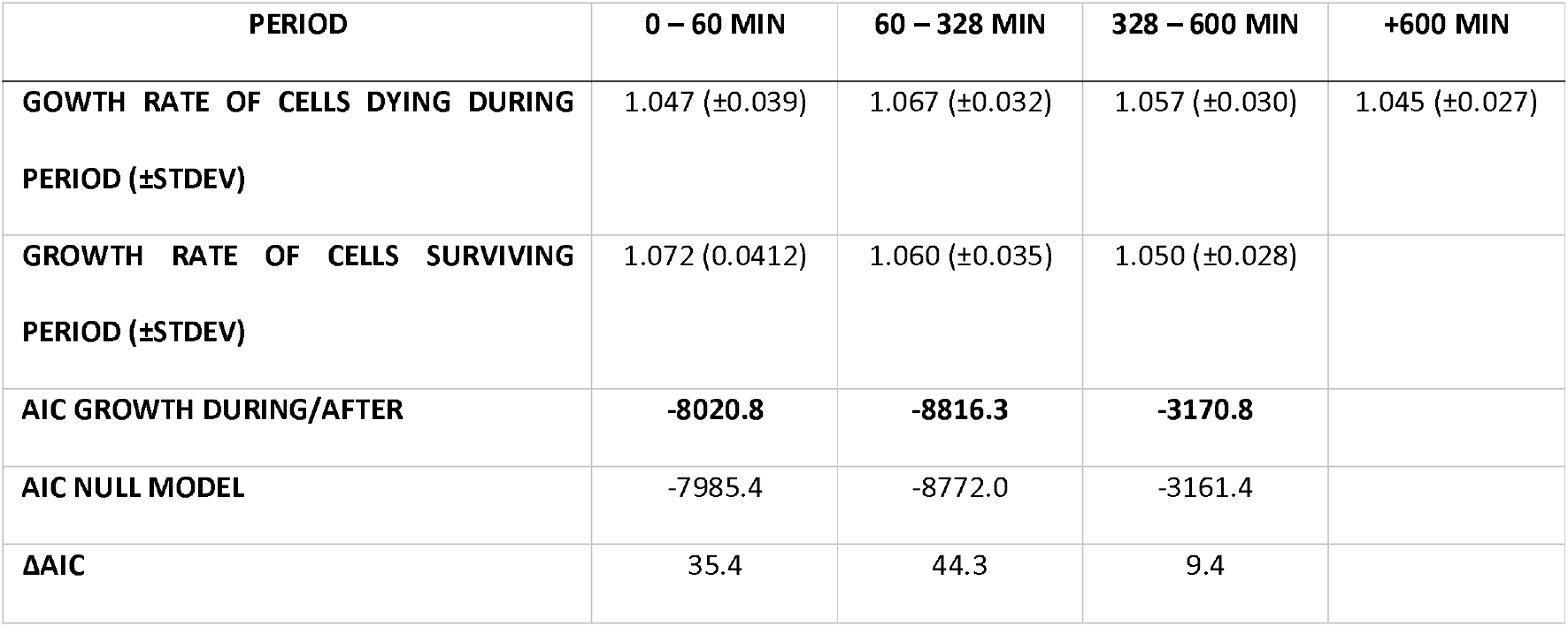
Differences in average growth rates of cells surviving past or dying during or after specific points in time. *Model comparison among a null model (intercept only model) and a model with the two groups of cells dying during a particular period or surviving that period as explanatory variable and the mean growth rate (per 4 min) as response variable. Three sets of models were fit, one for each period [0 min – 60 min; 60 min – 328 min; 328 min – 600 min]. A Gaussian error structure was used to fit these models.*

| PERIOD | 0 – 60 MIN | 60 – 328 MIN | 328 – 600 MIN | +600 MIN |
| --- | --- | --- | --- | --- |
| GROWTH RATE OF CELLS DYING DURING PERIOD (±STDEV) | 1.047 (±0.039) | 1.067 (±0.032) | 1.057 (±0.030) | 1.045 (±0.027) |
| GROWTH RATE OF CELLS SURVIVING PERIOD (±STDEV) | 1.072 (0.0412) | 1.060 (±0.035) | 1.050 (±0.028) |  |
| AIC GROWTH DURING/AFTER | **-8020.8** | **-8816.3** | **-3170.8** |  |
| AIC NULL MODEL | -7985.4 | -8772.0 | -3161.4 |  |
| ΔAIC | 35.4 | 44.3 | 9.4 |  |

### Robustness of growth rates within cells

In Fig. 2c, we investigated the average growth rate at a given time point. As cells can dynamically alter growth rates between the current and a future point in time, we correlated a cell’s current growth rate to its growth rate in the future, which provides a measure of robustness of growth. Overall correlation between an individual’s cell growth rate at a given time t and its growth rate in the immediate future (time *t*+1 [4 minutes later]) or future time points (time *t*+3 [12 min later]; time *t*+5 [20 minutes later]) was weak (Fig.4, Table 3), that is, growth rates are little robust and a cell’s current growth rate does not well predict its future growth rate. However, there was the exception for cells with intermediate growth rates (1.0-1.1). Cells growing in that range (Fig. 4, x-axis range 1.0-1.1) showed a strong correlation between current and future growth, illustrating robust constant growth for these cells. Growth arrested cells (growth rate <1 at time t) tended to increase in their growth rates at future points in time, while quickly-growing cells (growth rates >1.1 at time t) tended to decrease their growth rates (Fig. 4). Overall, we observed substantial temporal dynamics in growth rates with less robust growing cells (growth arrested and quickly-growing cells) being more affected by the concentration of antibiotics, which suggests that the growth rate responses to different concentrations of antibiotics shown in Fig. 2c are mainly influenced by the fastest growing and growth arrested cells.

**Table 3:** Model comparison among competing GAM models, for correlation of current growth rates to growth rates at future points in time. *Best supported model for each response variable (Growth t+1, t+3, t+5 respectively) is highlighted in bold. GAM models are fitted with the restricted maximum likelihood method, a smoothing parameter for Growth rate at time t with a shrinkage version of a cubic regression spline as smoothing parameter (bs=cs) and/or the concentration, including a model that accounts for the interaction of the two explanatory variables, growth rate and concentration of antibiotics. Growth rates are fitted with a Gaussian error structure and an inverse link function*.

|  | MODEL | DF | AIC |
| --- | --- | --- | --- |
| **GROWTH T+1** | **Concentr. + S(Growth by Concentr)** | **70.3** | **-441151** |
|  | Concentr. + S(Growth) | 17.0 | -440468 |
|  | S(Growth) | 11.0 | -438626 |
|  | Concentr. | 8.0 | -425891 |
| **GROWTH T+3** | **Concentr. + S(Growth by Concentr)** | **69.2** | **-421883** |
|  | Concentr. + S(Growth) | 17.0 | -421503 |
|  | S(Growth) | 11.0 | -420235 |
|  | Concentr. | 8.0 | -407188 |
| **GROWTH T+5** | **Concentr. + S(Growth by Concentr)** | **69.0** | **-412563** |
|  | Concentr. + S(Growth) | 17.0 | -412180 |
|  | S(Growth) | 11.0 | -410925 |
|  | Concentr. | 8.0 | -398709 |

### Population level responses

Much research investigating phenotypic resistance has been done in cell cultures, i.e. at the population level, which makes it challenging to partition mortality, reproduction and growth responses, characteristics that need single-cell level evaluation. Further, density dependence and resource depletion need consideration at the cell culture level but are controlled for by the microfluidics setup at the single-cell level. Scaling and comparing single-cell characteristics to population level patterns can therefore be challenging. In the following section we report on ‘control’ experiments done in cell cultures. For these cell culture experiments we took two approaches: CFUs were counted before and after each exposure period to antibiotics in order to mimic the recurrent application of antibiotics as in the single-cell experiments, or we evaluated cell densities (OD) under constant antibiotic concentrations to understand longer exposure to antibiotics. Cell cultures exposed up to the MIC showed similar qualitative dynamics in terms of numbers of CFUs when recurrently exposed to antibiotics (Fig. 5a). The higher the antibiotic concentration the stronger the reduction in CFUs, and the stronger the increase after relieving exposure to antibiotics, with dampened responses to the second and third exposure and subsequent releases. These patterns align with predictions of selection for phenotypic resistant cells as revealed for the single-cell findings. During the long recovery time after the second exposure, most cultures, except those previously exposed to 64 and 128 μg/ml, reached densities that likely imposed growth limitation due to resource limitation (i.e., the cultures approached stationary phase). The decline in CFU during the recurrent exposure shows the continued susceptibility of the cells after series of selection due to antibiotic exposure. When populations were permanently exposure to antibiotics (Fig. 5b), they, as expected, initial increased in density before densities were reduced to very low levels for all cultures exposed to MIC or supra-MIC concentrations, illustrating the susceptibility of the initial population and no occurrence of resistance mutations during the experiment.

**Fig. 5:**
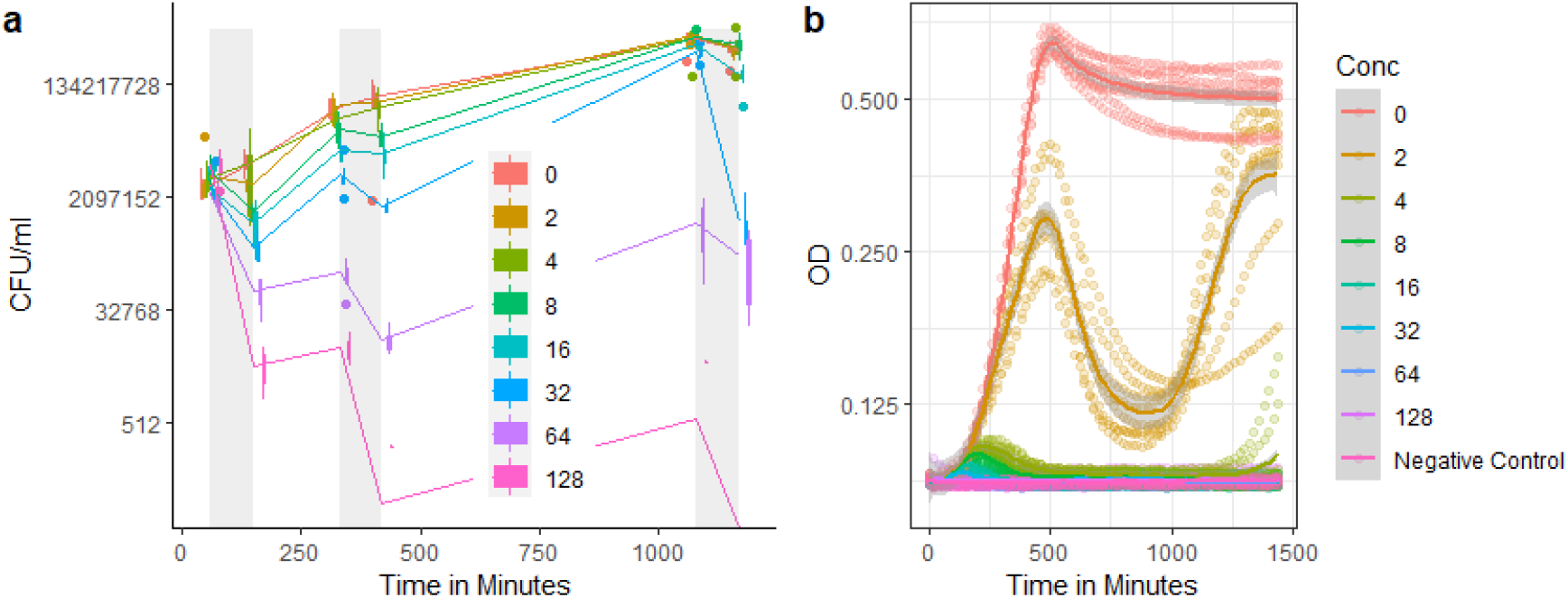
Colony forming units (CFU) (a), and optical density, OD_600_ (b) of cell cultures exposed to different levels of antibiotics across the duration of the experiment (a) or constant exposure to antibiotics throughout the experiment (b). The grey vertical bars (a) mark the exposure period to the antibiotics. CFU (a) are plotted as boxplots with line connecting mean number of CFU per time when CFU were counted. Single data points are outliers. Note, a jitter is added (a) for better visibility. For the OD curves (b) the dotted lines show the different replicates. the negative control (b) is no bacteria added to medium. CFU (a) and OD (b) are plotted on a log scale.

## DISCUSSION

We report on phenological characteristics of single-cells in *E. coli* in response to recurring exposure to a β-lactam antibiotic. Our findings suggest additional diversity of modes of phenotypic resistance, given that we show previously undescribed combination of characteristics. Compared to previous persister studies that quantify fractions of <0.1% persisters (2–4,8,10,25), we reveal a substantial larger fraction (~25%) of cells that survived 1.5 hours of antibiotic exposure, independent of the level of antibiotics (Fig. 1a, see also findings at cell culture for <64 μg/ml; Fig. 5a). This considerably larger fraction we find is best explained by the shorter exposure period to antibiotics we used to select for phenotypically resistant cells compared to the commonly used 3-5h exposure periods to select for persisters. We choose the exposure period as 1.5 hours exceed pharmacokinetics actions of antibiotics (46), illustrated for instance by the time to kill a cell by antibiotics often being comparable with bacterial generation time (16). For our conditions division time is much shorter than the 1.5 h exposure period and other studies report on substantially reduced mortality rates after 2h of exposure (16). Our single-cell survival curves (Fig. 1a) resemble those of a biphasic culture level killing curve, as is characteristic for persister cells. However the biphasic pattern cannot be directly compared, as the second phase of our findings concerns a higher fraction of cells and this phase might also be influenced by the release of the antibiotics after 1.5 h (as killing curves in other persister studies might be influenced by release time). Fractions of persisters varied among previous persister studies as selection conditions varied with respect to drug concentration, growth stage before selection (stationary exponential), preselection (priming of lineages, mutations), or growth media, to name a few. All of these conditions have been shown to alter persister fractions (2,3,15,29). In line with previous findings and as expected by selecting on persisters and against susceptible cells, the MIC marked a survival threshold, illustrating that the initial population was susceptible and not genetically resistant (Fig. 1), this susceptibility was also confirmed when evaluated at the cell culture level (Fig. 5). Our results show that the susceptible clonal parental population consists of phenotypically heterogeneous cells, whereby the cells have different capacities to sustain exposure to antibiotics, leaving predominantly phenotypic resistant cells after the first period of exposure.

We found reduced cell growth and division rates under antibiotic exposure, but this reduction was quantitatively different compared to most previously described persister cells. Growth reduction in our cells was about halved when exposed to antibiotics, while many other studies have reported growth arrest or near growth arrest of persister cells during antibiotic exposure (2,4,7,26). Single studies highlight that fast growth phenotypes with high efflux activity can avoid accumulation of antibiotics, findings that do not align well with the fact that antibiotics of exponentially growing cells accumulate within minutes in exposed cells (8,21). In contrast to previously described characteristics of persiter cells, after release from antibiotics, we did not observe resumption of normal growth within 2-4 hours (3,4). Our findings hence support studies that explored persisters derived from exponentially growing cells that showed longer lag times to regain growth, note these cells had been growth arrest during drug exposure, or highlighting substantial variation in lag times after antibiotic exposure release (10,25,47). A lack of growth resumption, as we found, is also a characteristic of persister type II cells and VBNC cells, however VBNC are found in complete growth arrest and are characterized by a small cell size, and persister type II cells are not environmentally induced but predetermined (18,26), features not found in our study and other studies (10). If classified, our phenotypic resistant cells fall best under specialized antibiotically induced persister cell definitions (9). Quantitative comparison to understand the resumed population growth at the cell culture level (Fig. 5) in light of the single cell growth reduction (Fig. 2) are challenging, as it remains opaque whether population growth dynamics are driven by altered cell division rates, altered cell growth, or altered mortality rates (48). In addition, with respect to heterogeneous selection and cell size, the average cell size remained constant over the course of the experiment (Fig. 2b) suggesting that selection did not favour small cells like VBNC cells or small birth sizes as described for persister cells (18,49), alternatively increased cell size should have been observed if increased frequencies in filamentation of cells occurred; here we only observed rare filamentation. Finally, variation in cell size did not increase substantially across the course of the experiment, which would have been predicted had higher proportions of small cells (VBNC) and filamentous cells simultaneously emerged, we also did not find other extreme morphological changes as reported for non-dormant persisters (10).

Our observations differ from genetically fixed resistant cells in various ways. First, at the single-cell level, unlike resistant cells, we did not find graded survival to different levels of antibiotics (Fig. 1 & 2a), which supports studies that show insensitivity of fractions of persisters up to ~100 μg/ml (15), although at the cell-culture level we observed graded responses in CFUs (Fig. 5). Second, our cells approximately halved their growth rates (~50% reduction in cell elongation rates) under drug exposure (Fig. 2c), whereas a 20% reduction in population growth (evaluated in cell cultures) is already considered to be an extreme growth cost for genetically fixed resistant cells (50). Third, in contrast to genetic resistant cells, our cells showed continued susceptibility to repeated exposure to antibiotics (Fig. 1, Fig. 5a). Finally, resistance is not expected for our single-cell findings, as our population of cells was directly derived from a susceptible, standard *E. coli* lab strain K-12 MG1655 (see also Fig. 5b). A heteroresistant population, i.e., one composed of multiple sub-populations that differ in their genetically fixed resistance to antibiotics (exemplified by different MIC), should have left, post exposure, a resistant population with graded responses to increased levels of antibiotic concentrations; we found none of these survival, division or growth rate patterns typical for resistant cells (Fig. 1&2) (50). Despite the isogeneic origin, our initial population was heterotolerant with a high fraction of cells (~25%) surviving the first exposure and continued showing dampened sensitivities with respect to mortality for following exposure periods (Fig. 2a). Such continued, but dampened, sensitivity could be explained by stochastic switching of smaller fractions of cells out of the tolerant state, which might be related to stochastic gene expression prior, during, and after exposure to antibiotics (51).

The growth characteristics of cells we describe in arising from exponential growing cells and not going into growth arrest have important implications for the evolution of genetic resistance. “Classical” persister cells only contribute to resistance development through repeated rounds of population expansion by their offspring, as they are growth arrested prior to and during antibiotic exposure, whereas our phenotypic resistant cells contribute before, during, and after antibiotic exposure to population expansion, and therefore provide continued opportunities for the accumulation of mutations towards resistance (31,37). Our phenotypic resistant cells cannot, a priori, be distinguished from susceptible cells by their reduced growth rate (26). Rare types of persister cells, that have previously described, did not arising from stationary phase cells and showed slow growth resumption post-exposure, these cells were almost growth arrested under antibiotic exposure (7,10), and we did not find such growth arrest for our cells.

Our finding that intermediate growing cells, that exhibited robust growth rates, had higher survival rates compared to growth arrested and quickly-growing cells (Fig. 3 & 4) is surprising to us, as we had predicted growth arrested cells to be the least affected by β-lactam exposure (2,4,14,26). Our findings might indicate that phenotypic resistant cells that exhibit high survival have more balanced and robust intrinsic metabolism (homeostasis) and less dynamic biochemical changes, which contrasts with molecular level findings that link phenotypic resistance to a perturbed biological network (33).

The phenotypic resistant state that our cells are characterized by is an induced state, and not an instantaneous switch, a pre-existing state (4), or a rare occasional stochastic switch to a persistence or tolerant state out of exponentially growing cells(26). Similar induced switching as we find has been found for different persistence (10,52) and aligns with findings where increased phenotypic resistance were induced by antimicrobial peptide exposure (7,53), though our cells seem not to instantaneously switch to their persistence stage, or at least require different lag times to do so. The emergence of our phenotypic resistant cells out of an exponentially growing population suggests that the evolution, or the maintenance, of phenotypic resistance is not conditional on nutrient limiting condition as has previously been suggested to be the favorable evolutionary route (54).

Our finding that cells cannot be easily identified prior to antibiotic exposure by their reduced growth rates also supports findings that there are various evolutionary routes to phenotypic resistance (4,5,9,10,33,55). This contrasts with our knowledge of the much better understood evolution of genetically fixed resistance, which tends to arise from mutations in a few defined genes that relate directly to the mechanism by which the antibiotic acts (33). The diversity of evolutionary paths to phenotypic resistance is illustrated by experimental evolution studies that evolved high phenotypic resistance by repeated exposure to ampicillin and showed that resistance was achieved through increased lag time (56); other experimental evolution studies, which exposed stationary phase cells, evolved phenotypic resistance without prolonged lag times (57), and still other experimental evolution studies found again other, lag time independent, mutations (33,58).

Future investigations should explore varying exposure time and varied inter exposure time periods to better understand how selection and characteristics depend on exposure periods. Prolonged treatment cycles can cause tolerant strains to attain resistance at higher rates (28), while multi-drug treatment can prolong time to persitence evolution (59), another dimension that could easily be explored in the future. Different strains should be explored to understand how specific the diverse phenotypic resistance mechanisms are, but such exploration should be systematic, as part of the diverse findings on phenotypic resistance might be caused by small differences in experimental settings not related to the main target (33). A rigorous phenomenological exploration, as we have done, can provide synergistic insights for tailored molecular exploration. Such molecular exploration will be interesting as of the expected diversity of molecular and evolutionary paths to phenotypic resistance. For instance, our cells did not show alterations in cell size, which suggests that our persistence might not be related to known resistance mechanisms to β-lactams, where filamentation rate is increased, which in turn stops ampicillin from efficiently targeting the cells (7,58,60). Our approach also illustrates the challenge and potential of scaling from single cell to the population level, a link that will provide a deeper understanding of the evolutionary potential towards resistance, but that is still rarely explored in detail (25). Our single-cell study is a cohort study and not a cross-cohort study like cell culture experiments, however cross-cohort studies can be achieved at the single cell level. Other differences, such as inter cellular communication, quorum sensing, nutrient limitation, or density dependence, among single-cell and cell culture explorations are more challenging to circumvent but these differences can reveal important conditions that help to reveal potential resistance mechanisms. The diversity and partly contrasting findings on phenotypic resistance characteristics, illustrates that we are still on the verge of understanding the role and mechanisms of such non-genetically determined resistance, and standardized and comparative studies are needed to solidify and advance our current understanding. Nevertheless, the importance of phenotypic resistance is reflected in our cell culture experiments, where population crashes were prevented by fractions of resistant cells when the population was exposed recurrently to antibiotics. Our results support the idea of diverse modes towards phenotypic resistance, where there is no pre-determined slow growth, no growth arrest during drug exposure, and no quick resumption of growth. These characteristics have implications for the evolution of resistance in providing evolutionary reservoirs and increased mutant supply; this combination of characteristics might even call for a wider definition of what persister cells are. As for applied implications on medical treatment, we need to be cautious about interpretation, because directly relating in vitro to in vivo results for antibiotic treatment remains challenging.

## Supporting information

Supplemental Information Statistics

## Acknowledgements

We thank Jens Rolff and Sophie Armitage for comments and discussions.

