## Supplemental Information Material and Methods for "Single-cell phenotypic characteristics of phenotypic resistance under recurring antibiotic exposure in *Escherichia coli*"

MATERIALS AND METHODS

**Bacteria cell culturing**

All experiments and bacteria preparation were done in filtered minimum growth media supplemented with 0.4% Glucose and 0.2% Casamino Acids (throughout called M9). One liter of M9- media consisted of: 11.28 g M9 Salts 5x, 2 g Casein hydrolysate, 980 ml destilled water, 100 µl 0.1 M CaCl 2, 200 µl 1 M MgSO4, 20 mg 20% glucose. To prepare the media that was used for the constant nutrition flow during the microfluidic experiments, we added propidium iodide (PI) at 1 µg/ml final concentration.

To derive the initial stock suspension for the microfluidic (single-cell) experiments, we incubated a small aliquot of a frozen bacterial stock of MG1655 overnight in 25 ml M9 in an incubator shaker (C24, New Brunswick Scientific) at 37°C and 200 rpm. The next day we inoculated 50 µl of the overnight bacteria batch culture in 2x35 ml fresh M9 media and incubated at 37°C for ~4 hours until the bacterial suspension reached an optical density (OD600) of 0.4-0.6 (corresponding to the most reproductive exponential growth phase). After these 4 hours we washed the bacteria by initially centrifuging at 4000 rpm for 10 minutes at 37°C, discarding the supernatant, resuspending in 500 µl of fresh media, transferring the resuspension to an Eppendorf tube, centrifuged that for 1 minute at 10000 rpm, discarded the supernatant, and resuspended the bacterial pellet in 200 µl.

**Microfluidic chip loading**

We then transferred the 200 µl of concentrated bacterial suspension to a 1ml syringe and injected it into the main (feeding) channel of a microfluidic chip (1). For the cells to enter the side (growth) channels of the microfluidic chip, we centrifuged the injected microchip at 400 rpm (~168g) for 10 minutes at 37°C. Subsequently, we checked the loading under the microscope and reinjected bacteria for a second time, this time without centrifuging, to ensure adequate loading (loading efficiency >85%). Next, we placed the microchip under an inverted microscope (Nikon Ti) with a temperature-controlled incubator (Oeko-Labs), connected the in- and outflow tubes to the chip, and media was flown at a constant rate of 300 µl per hour using a peristaltic pump throughout the experiment. Prior to connecting the tubes, we sterilized the pump tubing together with autoclaved rust-free connectors in a plastic zip bag by 15-minute exposure in a UV sterilizer. All experiments were conducted under these conditions at a stable temperature of 37°C and the single-cell experiments were conducted under constant flow of media at a rate of 300 µl per hour.

The microfluidic chip the cells grew in contained four main laminar flow channels (feeding channels) and hundreds of dead-end side channels, in the latter the rod-shaped bacteria-cells grew. Only one focal cell, the one closest to the dead-end of the side channel, was tracked for its growth (cell elongation), size, time of each division, and death. Cell death was determined by propidium iodide (PI) that emits a strong fluorescent signal when the cell wall lysis—the moment the PI can enter the cell—and bonds to the cells’ DNA. The cells loaded into the microfluidic chip came from an exponentially growing culture, and therefore the vast majority of cells loaded were in an exponential growth phase (2).

**Casting and mounting of the chip**

We used polydimethylsiloxane (PDMS) (Sylgard Silicone Elastomer Base and Curing Agent mixed in 10:1 ratio) to caste the microfluidic chips. Specially prepared molds produced with Sigatec SA Master (Smooth-Cast) were filled with PDMS and placed in an oven overnight (>12 hours) at 85°C. The next day we removed the casts from the molds under a clean air flow cabinet. The chip is designed with four main (feeding) channels, each holding 1200 side channels. Microfluidic inlets and outlets were punched by a sharpened gauge 22 needle at the start and end of each of the main (feeding) channels to later allow the flow of medium through the chip. Finally, we bonded the PDMS chip to a glass cover slide (24x60 mm) by treating the chip and slide for 30 seconds in an air plasma (plasma cleaner, PDC-002, Harrick Plasma), and immediately assembled the final microfluidic chip.

**Microchip setup and phenotypic resistance validation protocol**

To load cells into the chip we activated the chip surface by exposing it for 18 seconds to an air plasma, which makes the PDMS hydrophilic and allows injecting the media/20% PEG solution (400 µl M9- media + 100 µl PEG) inside the microchannels. We left the activated chip to incubate for at least 1 h before we loaded bacteria into the chip. PEG was used to prevent cell attachment to the chip and buildup of biofilms.

We flushed the in- and outflow connector tubes with media before connecting to the loaded chip, that was then mounted in a custom-designed microchip holder. The pump speed was set up to 300 µl per hour.

**Time-lapse Imaging**

Time-lapse images were taken at 4-minute intervals (15 frames/hour) and recorded by NIS Element AR software for data acquisition and microscope control. The fields of view were set up on each of the four main (feeding) channels of the chip, 15 fields of view per channel, together 60 for a whole microchip. Each field of view can record 23 dead-end-side (growth) channels with bacteria in them, that is, 1380 cells per chip could, theoretically, be tracked. However, the loading efficiency and the speed of the microscope table to change among field of views and autofocus does reduce this number. After defining the fields of view, two optical definitions (phase contrast and fluorescence) were set up. Fluorescence was used to detect cell death by PI emitting a red signal at cell lysis (PI binding to the DNA strand).

**Experimental treatment**

We chose antibiotic concentrations similar to previously explored ranges, that spanned from sub-MIC to >30 times MIC, and exposed cells to antibiotics for periods that would cover ~three divisions under non-antibiotic conditions. Post-exposure recovery times were also chosen on previous results reporting on regained growth of phenotypically resistant cells within 3-4 hours post-treatment (3–5). We choose ampicillin as antibiotic, a β-lactamase that inhibits bacterial cell wall synthesis*,* because this antibiotic has been extensively used in previous persister studies (3).

For the experiments with recurrent exposure (both single-cell and cell culture experiments), each experiment ran for 18 hours. The first exposure period was 60-150 minutes past onset of the experiments, the second exposure period was 330-420 minutes past onset of the experiments, i.e. 180 minutes past the first exposure period. The last exposure period was 1080-1170 minutes past onset of the experiments, i.e. 660 minutes past the second exposure period. When switching between treatment, that is between media (with or without antibiotics), we added a tiny air bubble to the tube, that allowed us to accurately mark the time of change of the media in the chip and prevented potential diffusion of the two media in the tubing. For the single-cell experiments, concentrations of ampicillin during the exposure periods were set to 2, 4, 16, 24, 32, 64 or 128µg/ml. As comparison and reference, data from another experiment was added as a control group, where cells were never exposed to antibiotics (1). Individual cell numbers tracked were 0 µg/ml n~898 (data added from other experiment (1)), 2 µg/ml n~334, 4 µg/ml n~337, 16 µg/ml n~669, 24 µg/ml n~330, 32µg/ml n~347, 64 µg/ml n~230, 128µg/ml n~351. Note, not for all measures was data on all cells available. For the experiments at the cell culture level, the concentrations of 0, 2, 4, 8, 16, 32, 64 or 128µg/ml were used, and in addition a negative control testing for potential contamination was done.

In addition to the single-cell experiments with 90-minute recurrent exposure periods we report on, we also ran two chips that had 30 minutes exposure periods. We do not report on these latter experiments as this short period did not lead to any antibiotic-exposure-triggered phenotypic effects and therefore are not presented.

**Image analysis**

After we collected the time-lapse images in ND format were exported to TIFF format and organized with customed programme TriFichier. The TIFF images were then analyzed with a custom written Visiopales programme (Opales Sarl, France). Afterwards, lab-written R scripts (6) checked for quality and errors were automatically corrected, see also (1). The final data matrix per experiment contained length of each cell at each 4-minute time point, fluorescence signal emitted from the PI to determine cell death (lysis) at each 4-minute time point, and time of division events of each cell throughout the experiment. From these matrixes we estimated in addition to size and division events, also time to death and growth rates.

**Statistical analyses**

*Survival analysis*

Kaplan-Meier survivorship curves (Fig. 1a) were computed with the survival package (survfit function; package [survival]) and compared among them with a Cox proportional hazard model in the R package [survival]. Cells that were still alive at the end of the experiment were noted to be right censored. Survival of 3496 cells were included in the analysis. A post-hoc test was then done to differentiate survival among the different concentrations of antibiotics by the pairwise_survdiff function of the survminer R package, applying a Holm p-value correction to account for multiple testing (for test see Supplementary file 1A). The probability of death curves (Fig. 1b) were plotted with ggplot2 package and a loess smoothing (geometric_smooth function; package ggplot2). Competing GAM models (Table 1) are fitted with the restricted maximum likelihood method (REML), for those models a smoothing parameter for age (time since the start of the experiment) with a shrinkage version of a cubic regression spline as smoothing parameter (bs=cs) and the exposure concentration was explored. A null model (intercept only model), a model with only one factor either age or concentration, and a model with an additive or an interactive effect among age and concentration was compared. A Gaussian error structure and an identity link function was used for these GAM models. Probability of death curves (Fig. 3b) are plotted with a loess smoothing function with 95% CI (geometric_smooth; package ggplot2).

*Growth rate analysis*

We excluded a few events where growth rates were very small (<0.8) or very large (>1.4) as such rates are biologically not feasible. Growth rate curves (Fig. 2c) are fitted with a GAM (with a loess geometric_smooth function). Model comparison was done by comparing AIC among competing GAM models estimated with restricted maximum likelihood method (REML) and a Gamma error structure with an inverse link function. For those models, a smoothing parameter for age (time since the start of the experiment) with a shrinkage version of a cubic regression spline as smoothing parameter (bs=cs) and the exposure concentration was explored. A null model (intercept only model), a model with only one factor either age or concentration, and a model with an additive or an interactive effect among age and concentration was compared. Note that the smoothing does weaken the actual more pronounced drop in growth rates post-antibiotic exposure, we still present the smoothed curves to be more consistent with the other analyses presented.

*Division rates*

Division rates (Fig.2a) are fitted with a loess smoothing function (geometric_smooth; package ggplot2) using a binomial error structure with a logit link function. In very rare cases of more than one division per 4-minute time step, that mainly happened when a filamentous cell divided into more than one cell at a time, we only considered this as a single division event to be able to use a binomial response variable (division yes/no) and an according error structure (binomial error structure and a logit link). Model comparison was done by comparing AIC among competing GAM models estimated with restricted maximum likelihood method (REML). For those models a smoothing parameter for age (time since the start of the experiment) with a shrinkage version of a cubic regression spline as smoothing parameter (bs=cs) and the exposure concentration was explored. A null model (intercept only model), a model with only one factor either age or concentration, and a model with an additive or an interactive effect among age and concentration was compared.

*Size*

Size curves (Fig. 2b) were fitted using a loess smoothing (geometric_smooth; package ggplot2) function. Model comparison was done by comparing AIC among competing GAM models estimated with restricted maximum likelihood method (REML). For those models a smoothing parameter for age (time since the start of the experiment) with a shrinkage version of a cubic regression spline as smoothing parameter (bs=cs) and the exposure concentration was explored. A null model (intercept only model), a model with only one factor either age or concentration, and a model with an additive or an interactive effect among age and concentration was compared. A Gaussian error structure and an identity link function was used for these GAM models. Note, smoothing functions (GAM) are highly sensitive at the extremes as partly illustrated by the diverse initial average cell sizes (Fig. 2b) of cells that came from the same parental population and that had not yet been exposed to antibiotics during the 60-minute lasting acclimation phase.

**Comparing mortality among different growth categories**

For cells exposed to concentrations at or above MIC we did a set of additional analyses. For these analyses we pooled cells across these concentration treatments.

*Growth before death*

We first tested whether cells that grew just before dying at high rates (>1.1), intermediate rates (1.1-1.0), or were growth arrested (including shrinkage) (<1) differed in their age at death. For these tests we used GLM’s with a gamma error structure and an inverse link function, age at death was the response variable and an intercept only model or a model that categorized cells into the three growth categories before death were compared using their AICs.

We then did a post-hoc test on the best model supported (GLM with Gamma error structure) to differentiate between pairs of growth categories in their age at death (Supplementary file 1B). For this, we used a Tukey test with the glht function in the mutlcomp R package.

*Robustness of growth rates*

To test for the robustness of growth rates we correlated growth rates at the current time *t* to the growth rate at time *t*+1, or *t*+3, or *t*+5. Curves (Fig. 4) were fitted using GAM models with the restricted maximum likelihood method, a smoothing parameter for Growth rate at time *t* with a shrinkage version of a cubic regression spline as smoothing parameter (bs=cs) and/or the concentration, including a model that accounts for the interaction of the two explanatory variables (growth at time *t* and time). Model comparison used the same GAM models with a Gaussian error structure and an identity link function.

**Movie of mother machine during experiment**

The time-lapse movie (Movie SI) shows one frame of view with 23 dead-end side channels. The focal cell is the cell most closely to the dead end (top of the channel). Below the side channels the main feeding channel with the laminar flow of media can be seen. Offspring cells are finally pushed out into this main feeding channel. The media also includes propidium iodide (PI), which emits at cell wall lysis (cell death) a red fluorescent signal. There recurring exposure of antibiotics is provided by the label.

**Cell culture experiments**

We inoculated 10 mL (1:100) from an overnight culture of *E. coli* MG1655 in the morning in fresh M9 medium and left cells to grow until the optical density of the culture at 600 nm was close to 0.2. At this point, we again inoculated the culture as before and added 200 µL to columns 1 to 8 of a 96-well plate. In the 9^th^ column, we added 200 µL of M9 medium as a negative control. We then incubated this initial plate for 1 h at 37°C and vigorous shaking (Supplemental Figure 1).

After this incubation time, we took a sample. For this, we transferred 50 µL of each column individually to a new multi-well plate, where the first column was completely empty and the rest of them until the column 6 contained 180 µL of NaCl for serial dilutions before we plated the samples on LB-agar plates. We used a total of eight multi-well plates for these serial dilutions, each plate for one column of the initial plate, and therefore eight LB-agar plates resulted from each sampling with 8 replicates (rows of plates).

Next, we exposed bacteria to ampicillin as for the single-cell experiments. From the initial plate, we transferred 100 µL of each well to a V-bottom-96-well plate containing 100 µL of serial dilutions of the antibiotic. The starting concentration at which bacteria were exposed was 128 µg/mL and the lowest, 2, passing through 64, 32, 16, 8 and 4 (note the single-cell experiments did not have 8 µg/mL exposure but 24 µg/mL). Each concentration was represented by a column in the multi-well plate, and every row constituted a replicate, obtaining a total of 8 replicates per concentration. The two last columns were always left as positive and negative controls, respectively.

Once we had prepared the plate, we incubated it for 90 min in the same conditions as above. When this exposure time passed, we took another sample as described before and plated a serial dilution on agar plates. We removed the antibiotic in the V-bottom-96-well plate by centrifuging 10 min at 4000 g and aspired the supernatant. We then resuspended cells in 200 µL of fresh M9 and incubated them for 3 h. After the recuperation time, we took another sample (serial dilution and plating on agar plates), and the bacteria were exposed again to decreasing concentrations of ampicillin as described before. The exposure time was again 90 min, after which we took the third sample, the antibiotic was again removed and a recovery period of 11 h in fresh medium followed.

Following these 11 h, we took a new sample, and transferred the bacteria to a new set of plates with decreasing antibiotic concentrations as we had previously done. We incubated one last time the cell cultures for 90 min. We then extracted the last sample after this final exposure time.

We incubated all the LB-agar plates at 30°C and counted the colonies (Colony Forming Units) after 16 h of incubation.

*Cell culture experiments constant exposure to antibiotics*

To track cell culture density at constant antibiotic exposure, the initial 8 replicates for each concentration of the antibiotic exposure were incubated at 37°C and shaking in a 96-well-plate and the optical density (OD600) was measured every 20 minutes for 24 h in a plate reader Synergy H1 (Biotek, Germany). Fig. 5b the OD curves are fitted with a loess smoothing function with 95% CI (geometric_smooth with a span of 0.1; package ggplot2).
