## Supplemental Information Statistics for "Single-cell phenotypic characteristics of phenotypic resistance under recurring antibiotic exposure in *Escherichia coli*"

Supplementary file 1

1A)

Table A: Post-hoc pairwise comparison testing survival curve differences among groups of cells exposed to the 8 different levels of antibiotic concentrations (Fig. 1a). Testing is based on a log rank test with p-value adjustment based on Holm. Cells that were alive at the end of the experiments are noted to be right censored. Values shown are pairwise p-values.

| **Antibiotics concentration** | **0** | **2** | **4** | **16** | **24** | **32** | **64** |
| --- | --- | --- | --- | --- | --- | --- | --- |
| **2** | <2e-16 | - | - | - | - | - | - |
| **4** | <2e-16 | <2e-16 | - | - | - | - | - |
| **16** | <2e-16 | <2e-16 | 1.000 | - | - | - | - |
| **24** | <2e-16 | <2e-16 | 0.668 | 0.335 | - | - | - |
| **32** | <2e-16 | <2e-16 | 1.000 | 1.000 | 1.000 | - | - |
| **64** | <2e-16 | <2e-16 | 0.054 | 0.013 | 1.000 | 0.372 | - |
| **128** | <2e-16 | <2e-16 | 0.668 | 0.331 | 1.000 | 1.000 | 1.000 |

1B)

Table B: Growth dependent mortality comparison

|  | **DF** | **AIC** |
| --- | --- | --- |
| **Age at death ~ Growth Category** | **4** | **22645.3** |
| **Age at death ~ 1** | 2 | 22671.1 |
| *Best supported model highlighted in bold. Age at death assigned to last growth rate. Only cells of groups exposed to >= 4 µg/ml included in analysis. Model comparison for GLM with Age at death as response variable, the growth categories (High, Mid, Shrinkage) as explanatory variable, or a null model (intercept only model), with a gamma error structure.* | | |

1C)

Table C: Pairwise comparison among different growth categories for age at death

| Pairwise comparison | Estimate | Std. Error | z value | Pr(>\|z\|) |
| --- | --- | --- | --- | --- |
| Mid - High | -0.0022818 | 0.0004944 | -4.615 | <0.001 |
| Shrink - High | -0.0007707 | 0.0006717 | -1.147 | 0.4801 |
| Shrink - Mid | 0.0015111 | 0.0005921 | 2.552 | 0.0281 |
| Adjusted p-values reported — single-step method | | | | |
