## Supplemental Fig. Methods for "Single-cell phenotypic characteristics of phenotypic resistance under recurring antibiotic exposure in *Escherichia coli*"

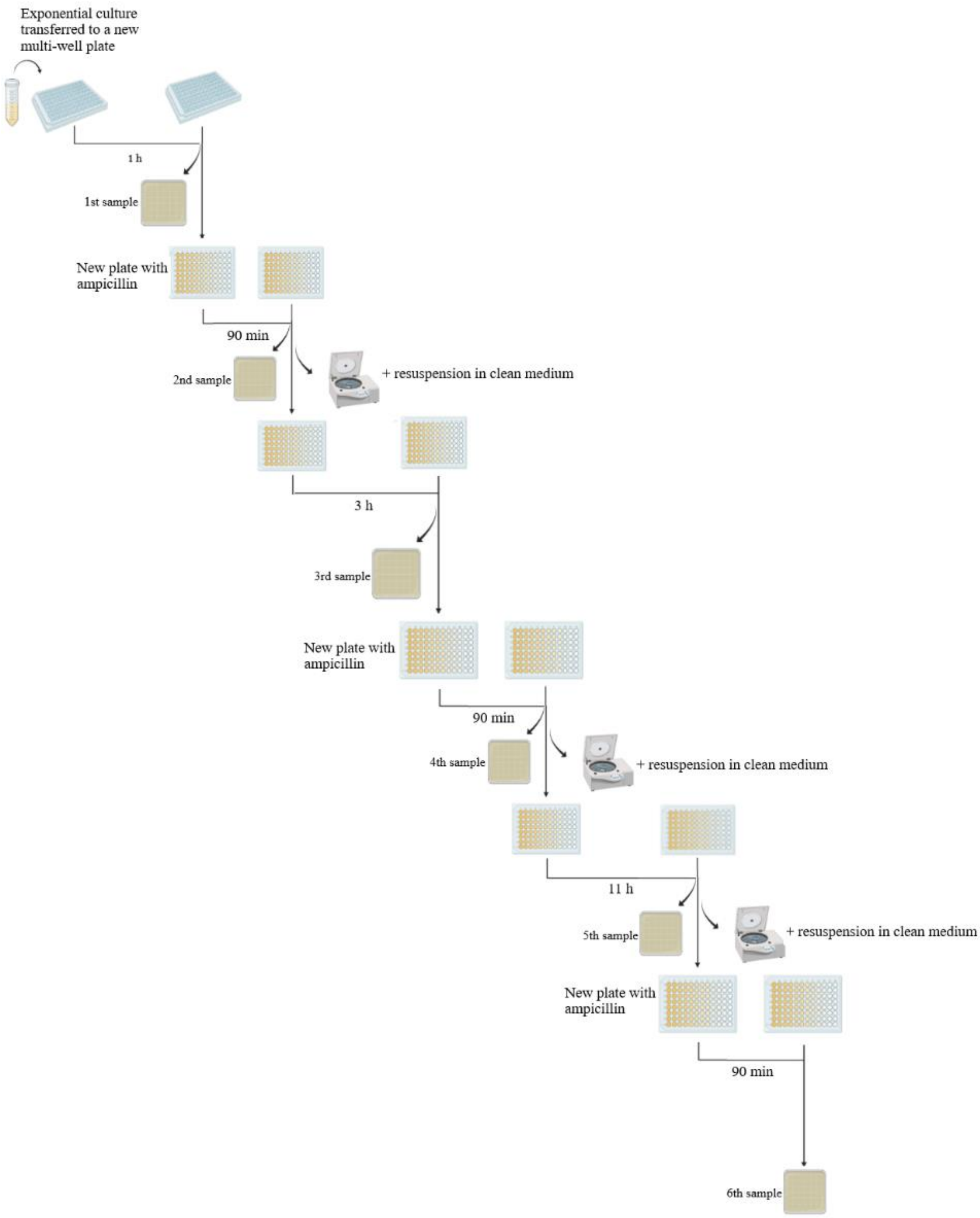


**Fig. S1: Workflow for the cell culture experiments.** The cell culture in exponential phase was added into a 96-multi-well plate and incubated for 1 h at 37°C and vigorous shaking. After this incubation time, a sample was taken from each well, serially diluted and plated in a LB-agar plate. Then, from the original multi-well plate, 100 µl were transferred to a new plate with the serial dilutions of the antibiotic, which was incubated again for 90 min. Each of the new plates with ampicillin that replaced the old one is highlighted in green. When the incubation time passed, another sample was taken, the plate was centrifuged to remove the antibiotic, cells were resuspended in new medium and a recovery period of 3 h followed. After this, a new sample was collected, the cells were again exposed to the antibiotic as described before and were left for incubation during 90 min. Once more, another sample was plated, and the antibiotic was removed to let the cells recover for 11 h. At the end, a new sample was collected, cells were exposed for the last time to the antibiotic for 90 min and the last sample was taken after this.
